# Stem cell-like memory T cells are closely related to follicular helper T lymphocytes

**DOI:** 10.1101/2024.06.01.596936

**Authors:** Nicolás Lalinde-Ruiz, David Santiago Padillo-Fino, Carlos Alberto Parra-López

## Abstract

Adoptive cell therapy has the potential to increase antitumor immunity by modifying cells *in-vitro* to expand lymphocytes that recognize and attack the tumor. The functional capacity and survival of the cells transferred to the patient heavily depends on the memory subpopulations that are being expanded in the laboratory, hence, obtaining early memory cells is desirable. The main objective of our work was to determine a strategy for *in-vitro* expansion of human stem cell-like memory T CD4 lymphocytes. Starting from naive cells, stimulated with a polyclonal agent supplemented with different combinations of cytokines from the common gamma family, we found that the combination of IL-7, IL-15 and IL-21 or IL-7 and IL-21 were the cocktails that produced a greater number of cells with a stem memory phenotype, measured by flow cytometry. Additionally, through the measurement of membrane proteins and *in-silico* analysis, a close relationship between stem cell-like memory and the follicular helper T cells differentiation program was established, which we believe contributes to a better understanding of the processes that underlie the generation and maintenance of memory and, therefore, may improve current strategies of expansion of T cells for immunotherapy purposes.

## Introduction

Within the strategies used in personalized medicine for cancer treatment, adoptive cell therapy (ACT) is one of the most promising types of immunotherapy modalities given the clinical responses reported in different clinical trials carried out in the United States, Europe, and China [1, 2]. To achieve an effective and lasting antitumoral response, the transferred cells must be able to expand immediately after transfusion and persist in circulation after the initial tumor reduction [2]. Both characteristics can be achieved by transferring early differentiated memory cells, which have a greater survival and antitumoral activity compared to their effector counterparts [3]. For this reason, the design of *in-vitro* strategies that allow obtaining high numbers of these cells has been subject to exhaustive research during the last decade.

IL-2 has been the most used cytokine for *in-vitro* expansion systems, generating large numbers of antigen-specific cells in a short time. However, recent evidence has shown that cells expanded with IL-2 not only display an effector phenotype, but there is also a significant percentage of these cells that are regulatory T cells (Tregs), which negatively affect the antitumoral capacity of ACT [4, 5]. Other cytokines belonging to the common gamma family such as IL-7 and IL-15, related to survival and homeostasis also have the ability to induce T cell proliferation without favoring Tregs expansion, and so their use in new experimental designs has been proposed [6, 7].

The identification of an intermediate memory population between naive and central memory cells, called “Stem cell-like” memory cells (Tscm), which can be expanded using IL-7 and IL-15, and whose potential of self-renovation and ability to differentiate in potent effector cells exceeds that central memory T cells, revolutionized *in-vitro* expansion systems for ACT [8–10].

IL-21, the last discovered cytokine belonging to the common gamma family, has proven to have the same capacity of inducing clonal expansion as IL-2, however it retains cells plasticity by inhibiting the expression of transcription factors associated with effector cells [11] and, recently, it was found to promote the expansion of the Tscm phenotype [12, 13].

So far, most Tscm expansion protocols have been focused on CD8 T cells [10] and experimental work leading the expansion of Tscm CD4 T lymphocytes in vitro is lacking, in this work, different in-vitro cell culture conditions were evaluated pursuing the expansion, and the expression of proteins associated with the response of Tscm CD4 T lymphocytes. Our results allowed us to establish that the combinations of IL-7 and IL-21 or IL-7, IL-15, and IL-21 are the best for Tscm cells expansion. That CD4 Tscm cells expanded under our culture conditions exhibited a pattern of protein expression and metabolic characteristics commonly associated with Tfh cells is intriguing. However, further studies are necessary to elucidate the interesting possibility that Tfh cells correspond to a differentiation status located in between CD4 Tscm cells and central memory CD4 T cells (Tcm) suggested by several results of our work.

## Materials y Methods

### Blood samples

This research was approved by an ethics committee within the framework of a project for identifying immunosenescence markers developed by the Translational Immunology and Medicine Group of Universidad Nacional de Colombia (No. 011-164-18 of June 15, 2018). Cells were obtained from blood units donated by the Instituto Distrital de Ciencia, Biotecnología e Innovación en Salud (IDCBIS), from ten healthy young donors between 18 and 30 years. PBMCS were isolated from blood units. Briefly, the content of the unit was transferred to 50 ml tubes and a series of saline solution washes were made in order to remove the anticoagulant. Subsequently, through density gradient separation with Ficoll® (Histopaque, Sigma Aldrich) PBMCs were obtained. Cells were counted, and their viability was evaluated by Trypan blue exclusion. Fresh cells were cultured, and remaining cells were cryopreserved in liquid nitrogen.

### Cell cultures

For cell cultures supplemented with different cytokines combinations, CD4 naive cells were enriched from fresh PBMCs. For the enrichment of this fraction, the EasyStep™ Human Naïve CD4+ T Cell isolation Kit II (Stemcell Technologies) was used according to the manufacturer’s instructions. After enrichment, cells were washed with AIM-V (Gibco, Thermofisher), and their viability was measured by Trypan blue exclusion.

For Tscm cells expansion, freshly isolated naive T cells were stimulated with magnetic beads coupled to anti-CD2, CD3 and CD28 antagonistic antibodies (Miltenyi Biotec) in a lymphocyte-bead ratio of 2:1. Cells were then cultured in 24 wells plates in 500 µl of AIM-V in a concentration of 2.5 x 10^5^ cells/well and the following combinations of cytokines (Cellgenix) were added: IL-2 low (10 U/ml); IL-2 mid (100 U/ml); IL-2 high (300 U/ml); IL-21 (10 ng/ml); IL-7 + IL-15 (10 ng/ml each); IL-7 + IL-21 (10 ng/ml each); IL-15 + IL-21 (10 ng/ml each); IL-7 + IL-15 + IL-21 (10 ng/ml each). In addition to this, there were two additional control treatments that corresponded to only beads and only cytokines without stimulation of CD3 and CD28. Cell cultures lasted 15 days and then cells were harvested for protein measurement by flow cytometry.

### Flow Cytometry

For flow cytometry, 5×10^5^ cells from each treatment were harvested and incubated with the respective antibodies for 20 minutes at 4°C. The following monoclonal antibodies were used: CD4-BV510 (Biolegend), CD45RA-APC-Cy7 (Biolegend), CCR7-FITC (Biolegend), CXCR5-PE-Cy7 (Biolegend), PD-1-PerCP-Cy5.5 (Biolegend), CD95-Alexa Fluor 647 (Biolegend), ICOS-PE (Biolegend), CD127-eFluor450 (eBioscience), and CD25-Alexa 700 (Biolegend). For membrane potential measurement, cells were labeled with MitoTracker™ RED CMXROS (ThermoFisher) prior to surface antibodies, as per the manufacturer’s instructions. Cells were acquired on a FACSAriaIIIu cytometer (BD Biosciences) and data was analyzed using FlowJo v10.0.7 software (Tresstar Inc.).

### Gen Set Enrichment Analysis (GSEA)

Transcriptome of the populations of interest (Tscm and Tfh subsets) was searched in the GSEA database. For Tfh, the full transcriptome code GSE130793 [14] was found sequenced with Illumina NextSeq 500. For memory subpopulations, the full transcriptome code GSE143215 [15] was found sequenced with Illumina HiSeq 2000. Transcriptome data for populations were downloaded and normalized according to the GSEA software. Subsequently, the pathways of interest were downloaded: STAT5 (M5947), STAT3 (M5897), Wnt-β-Catenin (M5895), Lef1 (M3572), Bcl-6 (M29904), NFAT (M2288), FAO (M5935), mTORC1 (M5924), ROS (M5938), Glycolysis (M5937). The normalized transcriptome and the corresponding pathway were the inputs of the software, enrichment graph, a heat map of the enriched genes and the values of each gene were the outputs of the software.

### Statistical Analysis

Statistical analysis was performed with Graphpad Prism v7 software. Comparisons between different times were made using Friedman’s repeated measures test with Dunn’s post-hoc analysis. For between-group analyses, Mann-Whitney and Kruskal-Wallis tests were used, as corresponded. Two-way ANOVA tests were carried out for analyzes of memory subpopulations between cultures. The significant differences are indicated in the graphs and the values were recorded in the legends of the figures.

## Results

### IL-7, IL-15, and IL-21 increase the expansion of Tscm lymphocytes

Most Tscm expansion protocols have focused on CD8 T cells [10], and very little information is available on this subset for the CD4 compartment. Considering that helper lymphocytes are responsible for regulating the generation and maintenance of memory and that, in their absence, the cytotoxic response is profoundly affected [16], in this work, we decided to focus on CD4 Tscm differentiation.

To evaluate the effect of different cytokines of the gamma-common chain in the expansion of CD4 Tscm cells, we magnetically enriched naïve CD4 T cells (> 95% purity (S1 Fig)), and then stimulated them with beads coated with anti-CD2/3/28 agonist antibodies in the presence of different concentrations of IL-2, and combinations of IL-7, IL-15, and IL-21. At day 15, the number of cells obtained, and their viability were measured by Trypan blue exclusion, and their memory phenotype was evaluated using flow cytometry (Fig 1).

**Figure 1.**
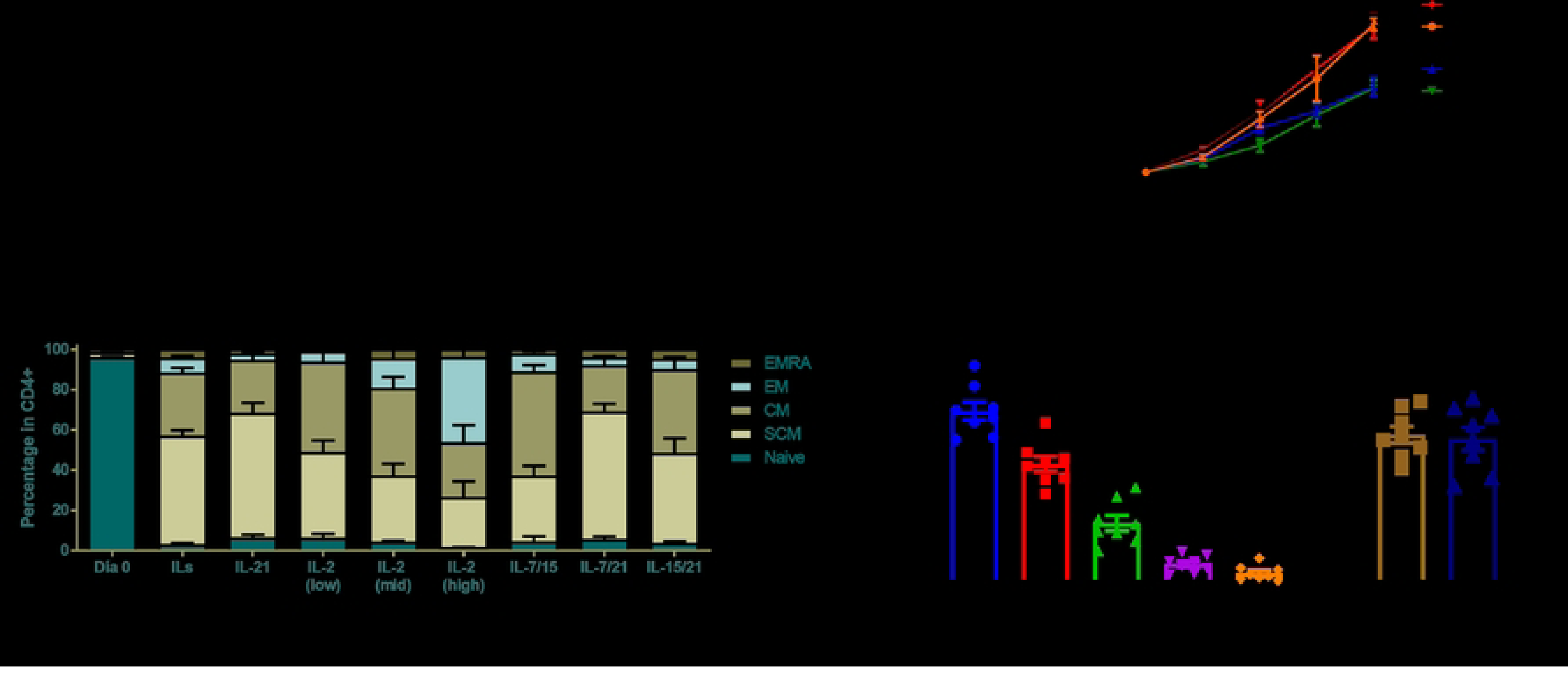
Effect of different cytokines of the common gamma family in the expansion of T helper cells. 2.5 x 10^5^ naïve CD4 lymphocytes were stimulated for 15 days with anti-CD2/CD3/CD28 beads in a 2:1 lymphocyte-bead ratio in the presence of **A.** Different concentrations of IL-2; 10 u/ml (low), 100 u/ml (mid), and 300 u/ml (high). **B.** IL-2 (100 u/ml); IL-7,15 and 21 (10 ng/ml each); only cytokines without stimulation beads (ILs); and only stimulation beads without cytokines (Beads). **C.** IL-7,15 and 21; IL-7 and 15; IL-15 and 21; IL-7 and 21; and IL-21. **D.** Percentage of memory subpopulations; Naïve, Stem Memory (SCM); Central memory (CM); Memory effectors (EM); Terminally differentiated (EMRA). **E.** Total numbers of Tscm cells at day 15. n=10. The data shown corresponds to the average + SEM. *<0.05; **<0.01; ***<0.005.

We first evaluated the effect that different concentrations of IL-2, the most common cytokine used for T cell expansion, had on the number of cells obtained at day 15. Using low (10 IU/ml), intermediate (100 IU/ml) and high (300 IU/ml) concentrations of IL-2, we observed that the culture with the intermediate concentration of this cytokine had a greater number of total cells (Fig 1A). Subsequently, we compared the expansion capacity of the intermediate concentration of IL-2 with the combination of IL-7, IL-15 and IL-21 (Fig 1B) and found that the use of these three cytokines together significantly increased the expansion of T cells at day 15, obtaining almost 2X the number of cells compared to IL-2 cultures.

We then aimed to evaluate the effect of different combinations of the three cytokines of our cocktail and found that all cultures supplemented with IL-15 had very similar numbers at day 15, however, when this cytokine was absent, numbers dropped close to those obtained with the intermediate concentration of IL-2 (Fig 1C). These results suggest that IL-15 is a cytokine capable of robustly expanding T lymphocytes, significantly outperforming IL-2. Similarly, it was observed that the combination of IL-15/IL-7 was sufficient to induce a high number of cells after 15 days of culture, and the addition of IL-21 apparently does not have a synergistic effect increasing the proliferation of stimulated T lymphocytes.

Although IL-21 did not induce the greatest expansion of total cells, its ability to induce an early differentiated memory phenotype, in contrast to IL-2, has been described [11]. Hence, we evaluated the cells for the presence of the following memory subsets at day 15 in our cultures: Naïve, Stem-like, Central Memory (Tcm), Effector Memory (Tem) and terminally differentiated effector (Temra) according to what was established by Gatinoni and Álvarez-Fernández [9, 12], who defined markers for the determination of Tscm cells (S1B and C Fig).

We found that the intermediate and high concentrations of IL-2 induced greater differentiation of the cells, observing a clear predominance of Tem cells, and a discrete population of Temra cells. Cultures with IL-7/15 and IL-15/21 had a higher frequency of Tcm cells, with a reduced presence of the effector memory subset, while the combination with IL-7/15/21, only IL-21 and IL-7/21, had a clear predominance of Tscm cells. Particularly, the culture with IL-7/21 was the one with the greatest expansion of cells with this phenotype, suggesting a synergistic effect between these two cytokines (Fig. 1D, S2 Fig).

Since percentages, being relative to the number of total cells, can be a biased measurement, the absolute number of memory stem cells was calculated (Fig 1E). Although cultures with IL-21 or IL-7/IL-21 were the ones that expanded the highest percentage of Tscm cells, the culture with the combination of IL-7, IL-15 and IL-21 had a greater number of both total and Tscm cells, suggesting that the use of these three cytokines together is necessary to expand this phenotype in large numbers.

### Tfh cells are induced by IL-21 correlate with Tscm cells

Since IL-21 has been canonically related to the Tfh phenotype, the observed effect of IL-21 in T cell differentiation led us to explore the induction of the Tfh phenotype under the proposed culture conditions of our work. To do this, Tfh cells were defined as CD4^+^CXCR5^+^PD-1^+^ [17], and its percentage was measured on day 15 in all cultures.

Confirming our hypothesis, the cultures with IL-21 were the ones with the highest percentage of the Tfh phenotype at day 15, being the cultures with only IL-21 and IL-7/IL-21 statistically different compared to those with intermediate and high concentrations of IL-2 (Fig 2A). Interestingly, the percentage of these cells was considerable (approximately 60%) in these two treatments, confirming the close relationship between IL-21 and this T cell subpopulation, and suggesting a possible role of IL-7 in their differentiation since the combination of these two cytokines showed a discretely higher percentage with less variation. Strikingly, cultures with IL-21 where IL-15 was also present, the frequency of Tfh was reduced, suggesting that IL-15 may be antagonizing the development of this particular helper subset.

**Figure 2.**
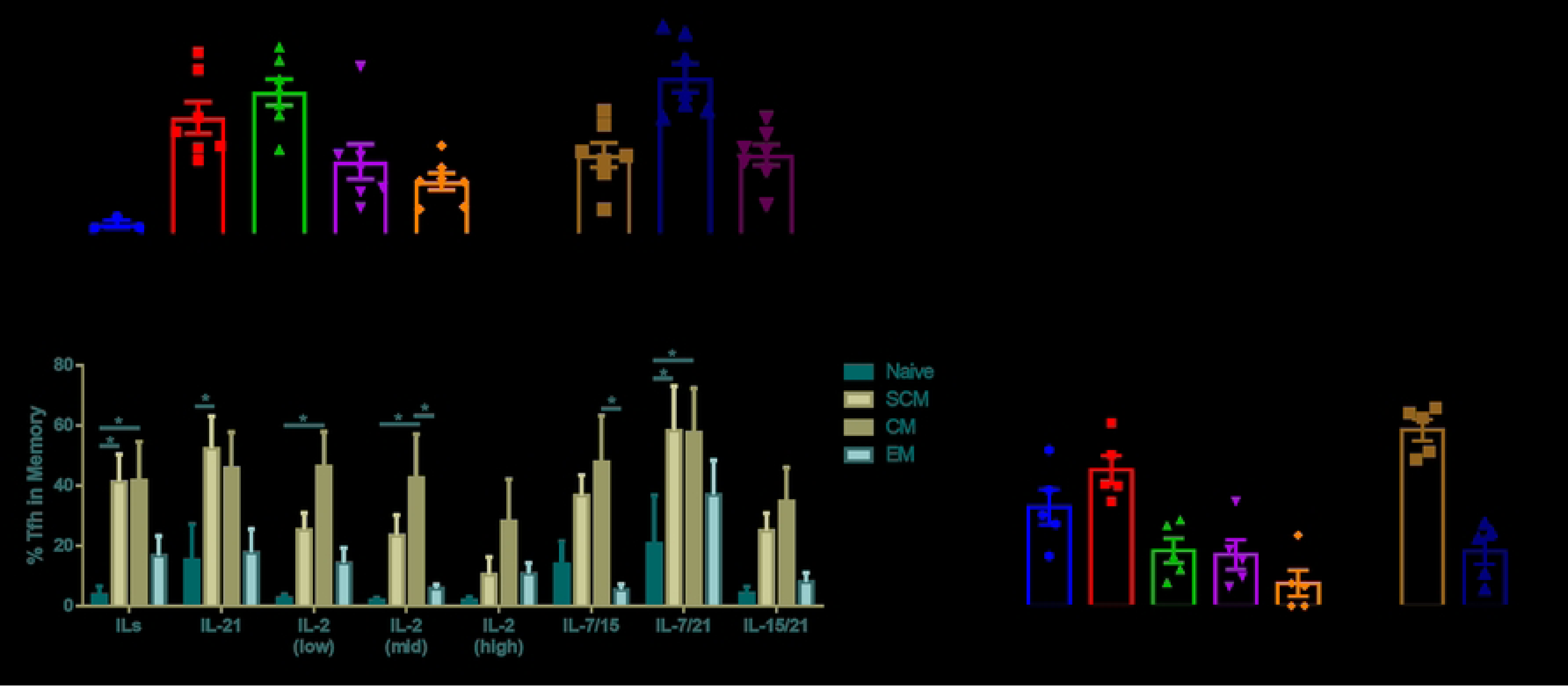
Expansion of CD4 follicular T helper cells on day 15 with different cytokines. 2.5 x10^5^ naïve CD4 T cells were stimulated for 15 days with agonist antibodies anti-CD2/CD3/CD28 beads in a 2:1 lymphocyte-bead ratio accompanied with IL-7, 15 and 21 (ILs); IL-21; IL-2 low (10 u/ml); IL-2 mid (100 u/ml) and IL-2 high (300 u/ml); IL-7 and 15, IL-7 and 21, IL-15 and 21. **A.** Percentage of Tfh cells (CXCR5^+^PD-1^+^) in total CD4 cells in the different cultures at day 15. **B.** Spearman’s correlation and p value between percentage of Tfh and Tscm cells on day 15 of the cultures. **C.** Percentage of Tfh cells in naïve cells; stem-like memory (SCM); Central memory (CM); Memory effectors (EM); and **D.** Comparison in Tscm cells. n = minimum 6. The bars correspond to the average +SEM. *<0.05; **<0.01.

The high percentage of Tfh cells in the cultures where there was also a significant percentage of Tscm lymphocytes suggested an association between these two differentiation pathways, probably induced by the effect of IL-21 and/or IL-7. To confirm this hypothesis, a correlation study was carried out between the percentage of both populations on day 15 of all cultures (Fig 2B). Here, we found that there was indeed a statistically significant positive association (r= 0.6614) between these two CD4 phenotypes. Given these results, we then sought to determine the percentage of Tfh cells within each of the memory subsets to see if all Tscm cells were also follicular helper cells and the how other memory subsets behave. For this, we evaluated the percentage of CXCR5^+^PD-1^+^ cells in naïve, stem-like memory, central memory, and effector memory (Fig 2C). This analysis was not runed in terminally differentiated cells due to the low numbers of this memory subset.

Memory phenotype of Tfh cells varied depending on the cytokines available in the culture. For instance, Tfh cells expanded with IL-2 had a predominately central memory phenotype, while those where IL-15 or IL-7 was present had both a central memory and a stem-like phenotype, and when only IL-21 was added, the majority of Tfh cells were also Tscm (Fig 2C). These results are coherent with our previous findings, which suggest that these two cytokines may have a synergistic effect in the follicular and stem-like cell differentiation. Subsequently, when comparing the percentage of Tfh cells within the Tscm between the cytokine regimens, it was clear that IL-21 and IL-7 promote both phenotypes, since almost 80% of all Tscm cells also expressed CXCR5 and PD-1 (Fig 2D). It is important to notice that once again, the addition of IL-15 was enough to reduce the percentage of Tfh markers in Tscm cells, although this was less noticeable than that observed with IL-2.

Once the expression of the Tfh markers was determined within the different memory subsets of the cells, the next step was to evaluate the overall memory phenotype of the Tfh cells expanded with the different gamma-common cytokines (Fig 3A). Interestingly, in almost all cultures, Tfh cells predominantly had a central memory phenotype, except in the culture supplemented with IL-2, where they had an effector memory phenotype, proving to be more differentiated cells. The cultures with both IL-7/IL-15, and IL-15/21 also showed a low percentage of Tfh cells with a Tscm phenotype, while all cultures in which IL-15 was absent had at least 20% of Tscm-Tfh cells.

**Figure 3.**
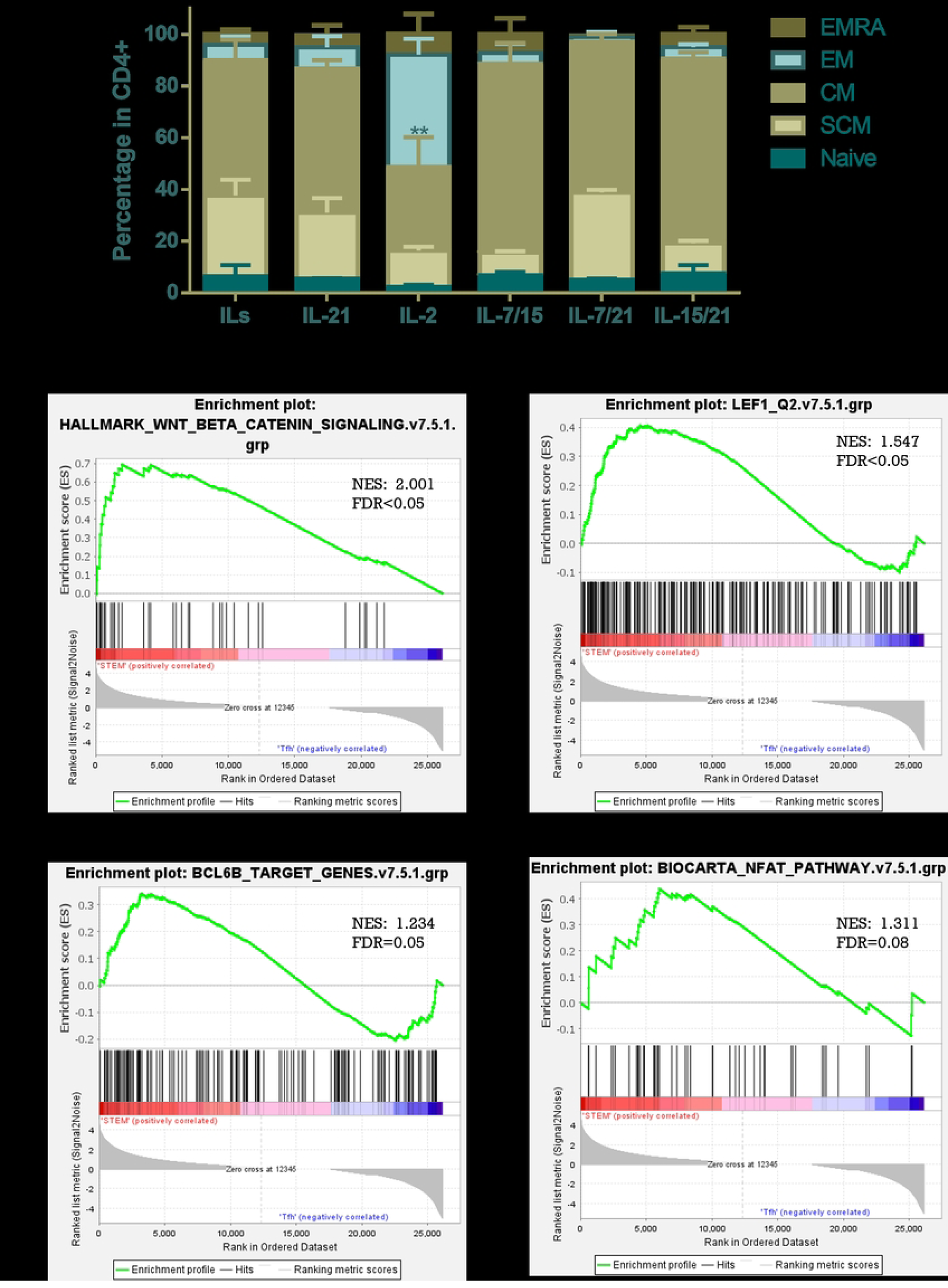
Tfh cells are more differentiated than Tscm cells but keep similar activation pathways. A. Memory profile of follicular helper CD4 LTs. 2.5 x10^5^ naïve CD4 LT were stimulated for 15 days with anti-CD2/CD3/CD28 beads in a 2:1 lymphocyte-bead ratio accompanied with IL-7, 15 and 21 (ILs); IL-21; IL-2 (100 U/ml); IL-7 and 15, IL-7 and 21, IL-15 and 21. Within the follicular helper CD4 LTs (CXCR5^+^PD-1^+^), the expression of CD45RA, CCR7 and CD95 was observed to define the memory subpopulations: Naive; Stem Memory (SCM); Central memory (CM); Memory effectors (EM); Effectors (EMRA). n=10. Bars correspond to average + SEM. Gene enrichment analysis of cellular self-renewal pathways. RNA-seq of Tscm (GSE143215) and Tfh (GSE130793) cells were obtained from public databases. Gene pathways were obtained from the GSEA page of previously published pathways. GSEA-reported gene set enrichment plots of **B.** Wnt-β-catenin (M5895), **C.** Lef1 (M3572), **D.** Bcl-6 (M29904), and **E.** NFAT (M2288) between Tscm (red) and Tfh (blue). The profile shows the continuous enrichment score (green curve) and the positions of gene set members (black vertical bars) in the rank-ordered list of differential gene expression. n= minimum 3.

### Tfh cells may originate and are closely related from Tscm cells

The previous findings suggest a close relation in the ontogeny of Tscm, Tcm and Tfh cells. In order to evaluate this potential convergence between the signaling programs induced by IL-21 and/or IL-7 in these three populations of CD4 T cells and had a better understanding of the origin of Tfh cells in the memory differentiation of helper cells, we decided to compare key intracellular pathways operating within these cells.

Self-renewal capacity of cells is one of the essential characteristics of Tscm cells and as they differentiate into Tcm or Tem this capacity is reduced [18]. Lef-1 and Tcf-1 are transcription factors that are involved with the pluripotent potential of Tscm, and both proteins are downstream of the Wnt-β-Catenin pathway [8, 10]. Taking into account that Tfh predominantly demonstrated a central memory phenotype, we compared the activity of the Wnt-β-Catenin and Lef1 pathways between single-cell RNA sequences available online for Tscm (GSE143215) and Tfh cells (GSE130793) [14, 15], using a Gene Enrichment Analysis (GSEA).

The vast majority of genes associated with both pathways were found on the left side of the graphs, showing a positive curve (NES>1.5), indicating a significant enrichment (FDR<0.05) of both signaling pathways for Tscm when compared to Tfh cells (Fig 3B and C). Particularly, protein genes associated with longevity and self-renewal such as Tcf7, Lef1, Notch1, Tp53, AMPD3 and PAK1 were found enriched (S1 File), confirming that Tscm cells have a much higher pluripotent potential than Tfh cells. When comparing Tfh and the other two memory populations: Tcm and Tem (S3 Fig), Tcm cells showed an enriched Wnt-β-Catenin and Lef1 pathways (2A and C Fig), in genes such as Notch1, Lef1, Tp53, Tcf7 and Wnt1, among others (S1 File). In contrast, it was observed that the Tfh had a greater enrichment of these pathways than Tem cells (S3 B and D Fig), which suggests that the Tfh, in terms of self-renewal capacity, are positioned between the Tcm and Tem subsets.

Bcl-6 is a key transcription factor in Tfh differentiation and is located downstream of the IL-21 receptor [17, 19]. Likewise, NFAT is another transcription factor that has been associated with Tfh, being responsible for the secretory profile of these cells compared to other Th subtypes [20]. For this reason, we evaluated the enrichment of these two transcription factors in Tscm cells with respect to Tfh (Fig 3D and E). Tscm cells showed an enrichment of genes associated with Bcl-6 and NFAT in comparison to Tfh cells. Of the Bcl-6 pathway, the enriched genes include STAT1, SOCS2 in Tscm, which have been associated with the effect of IL-21, as well as several genes associated with lipid metabolism such as LDLR and CISH (S1 File). Regarding NFAT, several genes related to this pathway (NFATc, PI3K and ligands) and cAMP-dependent kinases were found enriched in Tscm (S1 File). Comparing these pathways between Tfh and Tcm/Tem it was found that both Bcl-6 and NFAT were more enriched in the Tfh (S3E-H Fig), suggesting that these two pathways that have historically been associated with the Tfh phenotype are also play an important role in the differentiation of Tscm cells. Taking these results together, it seems that Tfh cells may originate from Tscm cells, and their activation pathways are kept throughout their differentiation, which seems to separate them from the other two memory subsets included (Tcm and Tem cells).

### Tscm and Tfh cells share a metabolic state compared to other memory subsets

Memory generation is a multifactorial phenomenon that includes increased cellular longevity [21], upregulation of key transcription factors [22], and epigenetic reprogramming of cells [23]. In the last decade, in addition to the aforementioned factors, the importance of metabolism in modeling the response and differentiation of immune cells has been explored [24]. Memory T cells must be able to quickly transition from a quiescent state to a highly proliferative and effector state, which requires a high energy supply from these cells [25]. In this sense, memory cells must have a high mitochondrial biomass capable of supplying the ATP requirements for protein production and cell division [26].

In addition to greater mitochondrial biomass, it is necessary for mitochondria to be able to produce high levels of ATP, so memory cells would be expected to have a high mitochondrial membrane potential (Ψm). For this reason, we evaluated the Ψm on day 15 of the cultures and determine the expression of memory (Fig 4A) and follicular helper markers (Fig 4B) in cells that had either a low or high Ψm.

**Figure 4.**
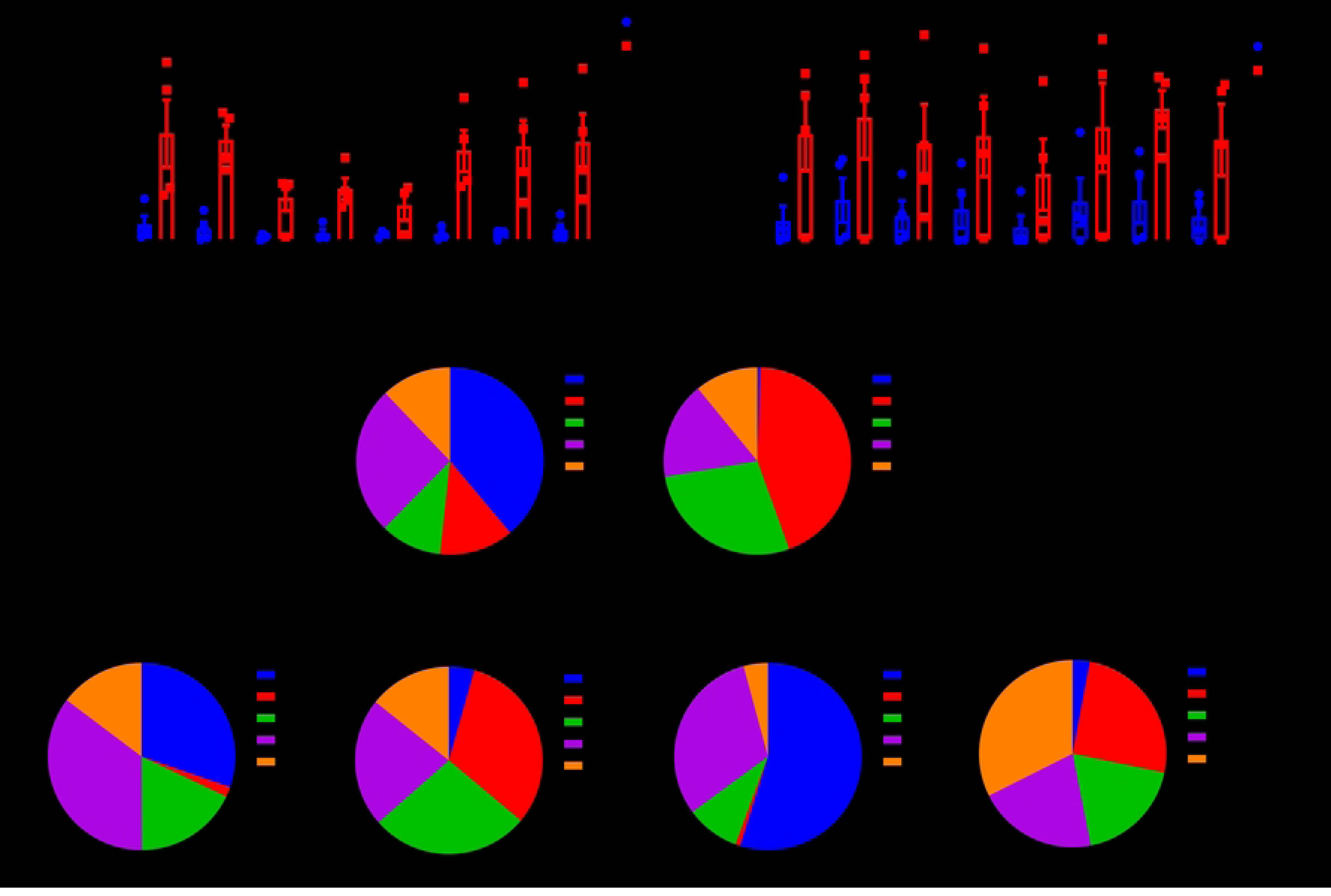
Mitochondrial membrane potential varies with the memory state of the cells. On day 15 of the respective cultures, the cells were harvested and stained with the MitoTracker CMX ROS probe. In the cytometry gate, **A.** Tscm and **B.** Tfh cells markers were evaluated in populations with low and high membrane potential. For (**C-E**), in the cytometry gate, the population with low membrane potential and the population with high membrane potential in total CD4 LTs was determined and within these gates the Naive memory phenotype was evaluated; Stem Memory (SCM); Central memory (CM); Memory effectors (EM) and terminally differentiated (EMRA) in cultures supplemented with **C.** IL-7, 15 and 21 (ILs), **D.** IL-21 and **E.** IL-2 mid (100 U/mL). n = 5. Two-way ANOVA. The bars correspond to the average +SEM.

Although no significant difference could be stablished, there was a clear tendency of expression of both helper subtypes markers in cells that had a higher Ψm. Of interest, in Tscm cells but not in Tfh cells, the addition of IL-2 diminished the expression of relative markers, which suggest that this cytokine may be modulating the metabolic state of mitochondria, favoring the production of ATP at the expense of Ψm. To further explore if the memory markers expression varied according to the Ψm state, we compared the percentage of the different memory subsets in cells with low and high Ψm in response to the different cytokines used for cell expansion.

The culture of IL-7, 15 and 21 was of particular interest, since this was the culture in which the greatest number of cells with Tscm phenotype was obtained, it was found that cells with a low mitochondrial potential had a high percentage of naïve cells (38.9%), followed by EM cells (25.6%) and that the rest of the populations were evenly distributed (approximately 12%). In contrast, cells with the highest potential corresponded to the less differentiated states of memory: SCM and CM (Fig 4C).

Additionally, this analysis was also done in the cultures with IL-21 and IL-2. These two cultures were chosen, since the inverse effect that these two cytokines have on metabolic aspects of the cells has been reported [13]. For IL-21, a fairly similar distribution of memory subpopulations was seen with respect to the combination of IL-7, 15 and 21, although a greater representation of effector memory could be seen in both low and high potential (Fig 4D). In contrast, in the IL-2 culture, the pattern of the memory phenotype in cells with a low membrane potential was maintained, while the distribution of the subpopulations changed dramatically in cells with high Ψm (Fig 4E). The phenotype with the greatest representation was the terminally differentiated cells, followed by EM. These findings suggest that IL-7, 15 and 21 induce metabolic changes in memory cells that allow them to have high mitochondrial membrane potential, while IL-2 seems to generate much more differentiated cells, with a high capacity for ATP production.

To further explore the similarity of the metabolic state of Tscm and Tfh cells, the membrane potential of the memory and Tfh populations was compared in the cultures with the combination of IL-7, 15 and 21, and in the intermediate concentration of IL-2. These two conditions were taken as an example, representing the conventional scheme used to expand LT and the one proposed as an alternative, both in recent studies [12] and in this work. In the culture with the combination of cytokines, in general, the Tscm and Tcm cells had a higher membrane potential, which was only surpassed by the Tfh (Fig 5A), while with IL-2 all populations seemed to have a higher membrane potential, except for the Tfh who remain the population with the highest mitochondrial membrane potential (Fig 5B).

**Figure 5.**
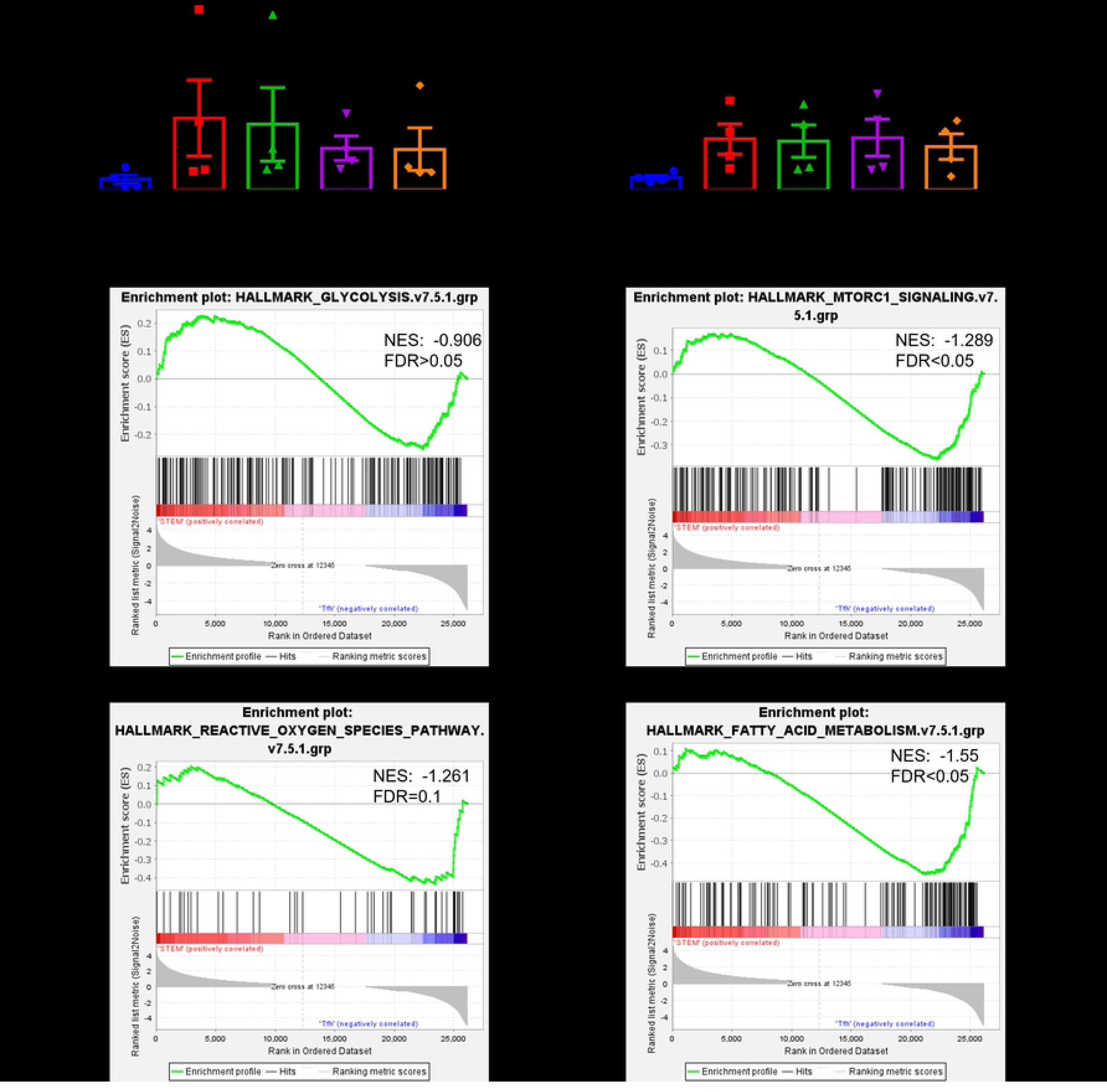
Tscm and Memory have a similar metabolic state. 2.5 x10^5^ naïve CD4 T cells were stimulated for 15 days with anti-CD2/CD3/CD28 beads in a 2:1 lymphocyte-bead ratio accompanied with **A.** IL-7, 15 and 21 (ILs) or **B.** IL −2 (100 U/mL) and stained with MitoTracker. n=5. Friedman test with Dunn correction. *<0.05. The bars correspond to the average +SEM. For (**C-F**) Gene enrichment analysis was performed with RNA-seq from Tscm (GSE143215) and Tfh (GSE130793) cells obtained from public databases and normalized for analysis by GSEA. Gene pathways were obtained from the GSEA page on previously published pathways. GSEA-reported gene set enrichment plots of **C.** mTORC1 (M5924), **D.** glycolysis (M5937), **E.** ROS (M5938), and **F.** FAO (M5935). The profile shows the continuous enrichment score (green curve) and the positions of gene set members (black vertical bars) in the rank-ordered list of differential gene expression. n= minimum 3.

As the Ψm observed for Tscm and Tfh was quite similar, contrasting with that of CM and EM cells, we runed a GSEA analysis of different key metabolic pathways. Canonically, memory populations have down-regulated pathways associated with glycolysis and, therefore, their glucose consumption is very low [27]. When comparing genes related to glycolysis and mTORC1, a master regulator of this metabolic pathway, both pathways were found to be enriched in Tfh, suggesting that these cells have a more effector profile and require a greater amount of energy (Fig 5C and D). Consistent with this, Tfh cells also showed a significant enrichment of ROS production and genes associated with this (Fig 5E), reaffirming that these cells are more differentiated and accumulate a greater amount of glycolysis byproducts. However, Tfh also showed upregulation of genes associated with FAO (Fig 5F), suggesting that this cell population uses both pathways for its energy requirements.

Although Tfh cells upregulate genes associated with glycolysis compared to Tscm cells, a negative association of these pathways was found, as well as the production of ROS in the Tfh compared to the CM and EM (S4 Fig), suggesting that these cells do not consume as much glucose as these other memory subpopulations and thus, prevent having a higher state of differentiation. Both Tcsm and Tfh also showed an upregulation of genes associated with FAO compared to the other memory subpopulations. Taking together, even though Tfh cells seem to have a stronger glycolic activity than Tscm cells, they heavily depend on fatty acid metabolism, unlike Tcm and Tem cells, which further validate our hypothesis that Tfh cells are closer to Tscm activity than to the others memory subsets.

### Automatized analysis confirms the closeness of Tscm and Tfh cells

The need for simultaneous assessment of multiple markers on cells analyzed by flow cytometry has prompted the development of fully automated methods that minimize the researcher’s subjectivity towards populations of interest (“gating”) [28]. Hence, we used a dimensionality reduction algorithm of the cells expanded using the different combinations of cytokines with the FlowSOM Software, and differences were found mainly in three of the eight subpopulations shown in figure 6. The first of them (Pop1) corresponded to a non-follicular (CXCR5^−^PD-1^−^) stem population (CD45RA^+^CCR7^+^CD95^+^) with an intermediate Ψm and an intermediate expression of CD25 and was found mainly favored in culture with IL-7/15/21, and the cultures with the lowest presence of this population were those of IL-2 and IL-7 and 15.

**Figure 6.**
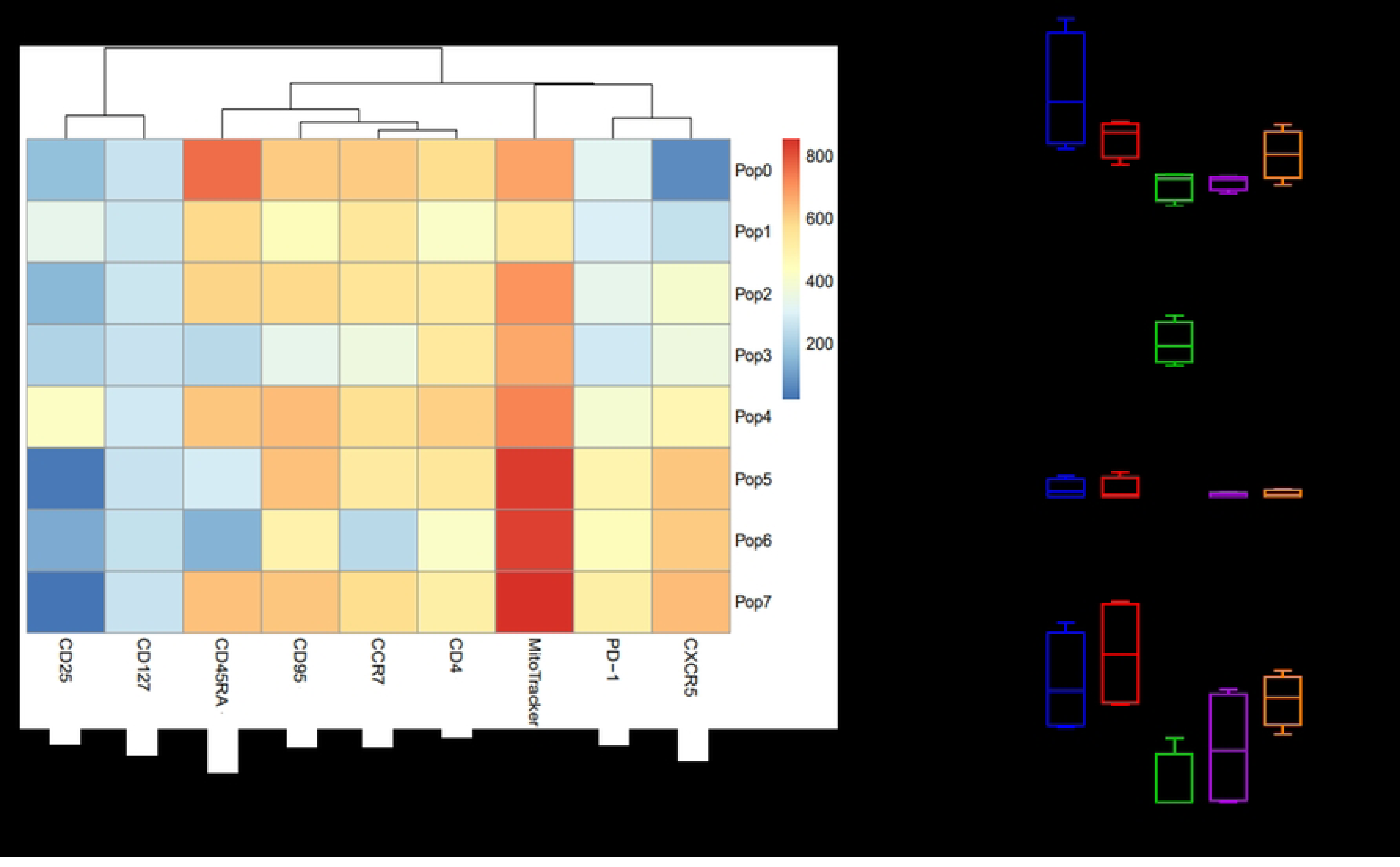
Automated cell clustering using FlowSOM software confirms the manual gating findings. A. Heat map showing the expression profile of each marker in the 8 groups identified in the concatenated file of the samples. **B.** Box-and-whisker plot showing the percentage of LTs of each group in the main groups identified in FlowSOM. n = 5. Data presented as means ± SEM. * <0.05.

Similar to this population, Pop7 was also significantly decreased in IL-2 and IL-7 and 15 cultures, although a lot of variability was seen in the latter. This cluster was a population of Tscm cells with a Tfh phenotype and high mitochondrial potential. Finally, Pop3, which unlike the other two was predominant in cultures with IL-2, was a population with an EM phenotype. The results obtained with the automated analyzes confirm that Tscm cells either with a Tfh (pop7) or non-Tfh (pop1) are enriched when stimulated with IL-7 and 21, and that Tem cells are favoured when adding IL-2. These findings were similar to those obtained with our manual gating strategy, which increases the robustness of the conclusions reached in this study.

## Discussion

ACT is one of the most promising immunotherapy modalities for cancer treatment due to the clinical responses that have been reported in different clinical trials carried out mainly in the United States, Europe and China [2]. To achieve an effective and durable antitumor response, the transferred cells must be able to expand immediately after transfusion and persist in circulation subsequent to initial tumor control [29]. Both characteristics can be achieved by enriching early differentiated memory cells i.e., SCM and CM, as these have greater survival and antitumor activity than their effector counterparts [3]. For this reason, the search for *in-vitro* strategies that allow obtaining high numbers of these cells has been the subject of exhaustive research during the last decades.

This work sought to identify culture conditions that would allow the expansion of Tscm cells. To do this, eight different culture conditions were tested, and it was evaluated which combination and concentration of cytokines was the most appropriate to obtain a large number of this memory subpopulation. Originally, cultures were started from total PBMCs (data not shown), which were stimulated with a polyclonal agent and supplemented with the corresponding cytokines. Regardless of the treatment, it was not possible to maintain a sustained expansion of the cells, observing that the vast majority had an effector memory phenotype, which was accompanied by a decline in viability after day seven of cultures. For this reason, we proceeded to enrich the naive fraction to start the next cultures, since these cells are capable of differentiating into all other memory subpopulations [30].

Under these new starting conditions, in all cultures a progressive growth of the stimulated cells could be observed, maintaining a minimum viability of 80% during the 15 days of culture. Coherent to published literature [6, 12], the IL-7/15, IL-15/21 or IL-7/15/21 cocktails significantly exceeded what was observed with any of the other five culture conditions (Fig 1). In the absence of IL-15, IL-7 and 21 induced an expansion very similar to that observed with IL-2, which suggests the high capacity of IL-15 to promote lymphocyte proliferation compared to other cytokines of the common gamma family.

Classically, IL-2 has been the cytokine that has been associated with T cell proliferation [31], but as we show in our results, IL-15 generated an increase 1.6 times greater. In addition to sharing the common gamma receptor, IL-2 and IL-15 receptors also have the beta subunit (CD122) in common, so the differential expansion caused by these cytokines is expected to depend on signaling to through the alpha subunit of its receptors [31]. The alpha subunit of the IL-15 receptor is commonly not found in T cells but in APCs [7], suggesting that signaling downstream of CD25 may downregulate the proliferative capacity of T cells, compared to the action of CD122 and CD132 alone.

The main objective of this work was not limited to the large-scale expansion of cells, but also sought to achieve the induction of an early differentiated memory phenotype i.e., Tscm. After counting and measuring their viability, the memory phenotype of T lymphocytes was assessed by flow cytometry at day 15 of the cultures. The addition of CD95 to canonical memory markers (CD45RA and CCR7) allows Tscm cells to be differentiated from naïve cells [9].

Using these three markers, it was found that both in percentage and in absolute number the cultures that expanded the most Tscm cells were IL-7/15/21, IL-7 and 21 and only IL-21 (Fig 1 and S2 Fig). Similar to the effect seen with IL-15 in terms of expansion, IL-21 was the cytokine that greater induced the stem-like phenotype, since in its absence the percentage of these cells was limited. Interestingly, the culture of IL-15 and 21 had a predominant Tcm phenotype, considerably reducing the percentage of Tscm cells. In absolute numbers, this difference is not so clear due to a high variability between the replicates, but it can also be explained by the high expansion promoted by IL-15. In this sense, it could be hypothesized that IL-15 is more similar to IL-2, being mainly responsible for promoting high lymphocyte proliferation, while IL-7 and 21 would be more related to maintaining a poorly differentiated memory phenotype, although this effect is clearer for IL-21 than for IL-7.

Based on these results, it is possible to conclude that the most appropriate for T cell expansion protocols is the combination of IL-7, 15 and 21, since it results in a considerable increase of the initial cells number (16x), of which approximately 50% have an Tscm phenotype, and the remaining percentage corresponds mainly to Tcm cells, which, although they are a little more differentiated, still have good antitumor activity and much longer survival than effector cells [32]. Another alternative is the combination of IL-7 and 21, with which a fairly close absolute number of Tscm lymphocytes was obtained. This treatment is proposed as an alternative because it has been observed that the less differentiated the transferred cells are, the smaller number of cells is also needed to obtain a favorable clinical response [9], and the use of two cytokines reduces costs, which is a factor that must be taken into account when proposing experimental strategies for clinical use.

CD4 T cells are highly plastic cells and depending on their cytokine production they are classified into Th1, Th2, Tregs, Tfh cells, etc [33]. For example, IL-2 was originally thought to be a Th1 cytokine, but is now considered to be secreted by T cells before they commit to either lineage [21]. On the other hand, IL-21 has been strongly associated to the Tfh phenotype and its function was originally described in NK and B cells [34]. According to our results, IL-21 is capable of inducing the generation of early differentiated memory cells. Being involved in these two phenotypes of the helper response, we wanted to explore the relationship that both differentiation programs had in order to better understand the mechanisms of memory generation and maintenance.

In a similar way to the percentages of Tscm cells in the cultures with IL-21, except in that of IL-15 and 21, we found the highest frequency of Tfh cells, and a positive correlation could be established between these two populations (Fig 2), showing that there is a close relationship between these two phenotypes. To explore if this was the same subset of cells, the expression of the proteins used to define Tfh cells (CXCR5, PD-1) was evaluated in the different memory subpopulations, as well as the memory profile of the overall Tfh cells.

We found that Tscm and Tcm cells were the subsets with the highest expression of the Tfh markers, ranging between 20 and 60%. In cultures supplemented with IL-2, especially at a high concentration, the co-expression of CXCR5 and PD-1 was not as high and was restricted to central memory cells. The culture with the highest percentage of Tfh in Tscm and Tcm was observed with the combination of IL-7 and 21, followed by only IL-21 (Fig 2A). This observation follows the pattern that has been identified so far, where IL-7 and 21 seem to have a synergistic action promoting an Tscm memory phenotype and now a Tfh phenotype, suggesting very similar signaling pathways.

In contrast, IL-15, although it does not seem to antagonize the follicular differentiation program as drastically as IL-2, it does seem to separate itself from IL-7 and 21. In this sense, a gene enrichment analysis was performed of two pathways downstream of the gamma-common family of cytokine receptors: STAT3 and STAT5. A pattern was found where Tfh and Tscm cells showed enrichment of STAT3, while Tem favored STAT5 (S3 Fig). In the literature, it has already been documented that STAT5 antagonizes the Tfh program and favors Th1 cells [35], so it can be thought that both IL-2 and IL-15 signal preferentially through STAT5, inducing a phenotype more differentiated, while IL-7 and IL-21 promote a Tscm/Tfh phenotype through the activation of STAT3 and to a lesser extent STAT1 (S1 File).

On the other hand, when the memory phenotype of the Tfh cells was determined, we found that they were mostly central memory cells, except when IL-2 was added, where the cells were predominantly Tem (Fig 3A). However, IL-21 induced a significant percentage of Tfh-Tscm cells. It seems important to us to emphasize within our findings this percentage of Tfh cells that maintain the expression of CD45RA and CCR7, since in several works that have studied this CD4 subset, they exclude “naive” cells in their gating strategies based only on these two markers [17, 36, 37].

The predominance of a Tcm phenotype in the bulk of Tfh cells suggests that these cells are more differentiated than Tscm cells. To evaluate this hypothesis, a GSEA analysis of pathways associated with self-renewal, an essential characteristic of stem cells, was performed. Indeed, Tscm cells showed greater enrichment of the Wnt-β-Catenin and Lef1 pathways than Tfh cells (Fig 3B and C). Likewise, the Tcm also had a greater enrichment of these pathways than the Tfh, who only showed a positive NES value compared to the Tem (S3 Fig, S1 File).

In addition to what has been evaluated so far, the metabolic state of the cells is another factor that must be taken into consideration when evaluating *in-vitro* expansion strategies, since this is also determinant for memory generation and cell exhaustion [25, 38]. Mitochondrial membrane potential is a good indicator of the ability of cells to produce ATP in order to sustain their effector functions [26]. Tscm cells have been reported to have a high membrane potential compared to naive cells, which contributes to the ability of these cells to respond quickly to antigen stimulation. Naive cells, in contrast, require *de-novo* mitochondrial biogenesis to be able to supply the energy requirements for proliferation and cytokine production [39].

Under our culture conditions, we found that in the presence IL-2, the cells that had the highest membrane potential were the EMRA cells, which in these cells, rather than talking about their ability to produce large amounts of ATP, has been related to an uncoupling of mitochondrial function with the energy needs of the cell, which is associated with an induction of apoptosis intrinsic pathway (Fig 4). This suggest that Tem and TEMRA cells are not good candidates for ACT as they are very close to apoptosis.

In contrast, the cultures supplemented with IL-7, 15 and 21 and only IL-21, which induced the higher percentage of Tscm cells, were also the cultures where these cells had the greatest mitochondrial potential. This shows the close relationship that exists between metabolic characteristics i.e., mitochondrial status and the generation of early differentiated memory cells. Similarly, the mitochondrial potential of Tfh was also evaluated independently and it was found that, similar to Tscm lymphocytes, these cells have a high mitochondrial membrane potential, far surpassing naive cells.

Canonically, immunometabolism studies have raised a dichotomy between memory and effector cells, where the former predominantly use a catabolic metabolism based on the oxidation of fatty acids, and the latter an anabolic metabolism based on aerobic glycolysis [27]. Following this theory, the upregulation of genes associated with glycolysis and FAO between Tfh and Tscm was compared and interestingly it was found that both pathways were upregulated in Tfh (Fig 5). When comparing Tfh with the other two memory subpopulations, it was found that Tfh upregulate proteins associated with FAO, while Tcm and Tem cells have a positive association with glycolysis and ROS production compared to Tfh.

The high mitochondrial potential of Tfh, as well as its greater FAO capacity, could be related to a high energy need, which cannot be met solely by the consumption of glucose, since being found in germinal centers, the B cells that are proliferating create a limitation of resources, making these cells favor a metabolism based on the catabolism of fatty acids, which could be induced by signaling through PD-1, which is one of the flagship markers of this cell population and is downstream of NFAT [40], another Tfh-related transcription factor that was also found upregulated in this population (Fig 2). The results found on the metabolism of Tfh in relation to memory subpopulations favor the hypothesis that these are closer to Tscm than Tcm, by favoring FAO over glycolysis.

Manual findings using conventional gating strategies were corroborated using an automated software capable of classifying samples into clusters in an unsupervised manner in order to reduce the subjectivity of the analyzes [41, 42]. Using this approach, the expansion of Tscm cells with a Tfh phenotype could be confirmed (Fig 6), so it can be concluded that these two populations do indeed have a close relationship and the pathways that have been established for follicular induction must be taken into account in new Tscm expansion strategies.

Based on the results presented here, it is possible to propose that Tfh cells comprise an intermediate population between the Tscm and Tcm state and rather than representing an analogue to the other Th subtypes, follicular helper cells could be thought of as a state of memory differentiation, which can latter differentiate into any other phenotype [33], which could explain the very high plasticity and capacity to produce cytokines associated with the other Th subtypes that this CD4 subset has [43]. However, it is important to continue generating more experimental evidence to support this theory.

Finally, we also highlight from this work the study of Tscm in the helper compartment, since the vast majority of works on this memory subpopulation have focused on CD8 T cells [15]. In the context of ACT, the study of CD8 lymphocytes has been favored since they are cytotoxic cells capable of directly attacking the tumor. However, CD4 T cells, in addition to having a potential cytotoxic capacity, they are also responsible for modulating the entire immune response, including CD8 T cells, B cells and macrophages, so their antitumor potential may be much more beneficial [44].

## Acknowledgments

The authors express their gratitude to Instituto Distrital de Ciencia, Biotecnología e Innovación en Salud (IDCBIS), as well as the volunteers that generously donate their blood, which made this research possible. This study was supported by funding form the Universidad Nacional de Colombia (Hermes 57494 and 57457) and from MinCiencias (920 grant, RC 800 of 2023; 841 grant, RC 903 of 2019).

## Supporting information

**S1 Figure.**
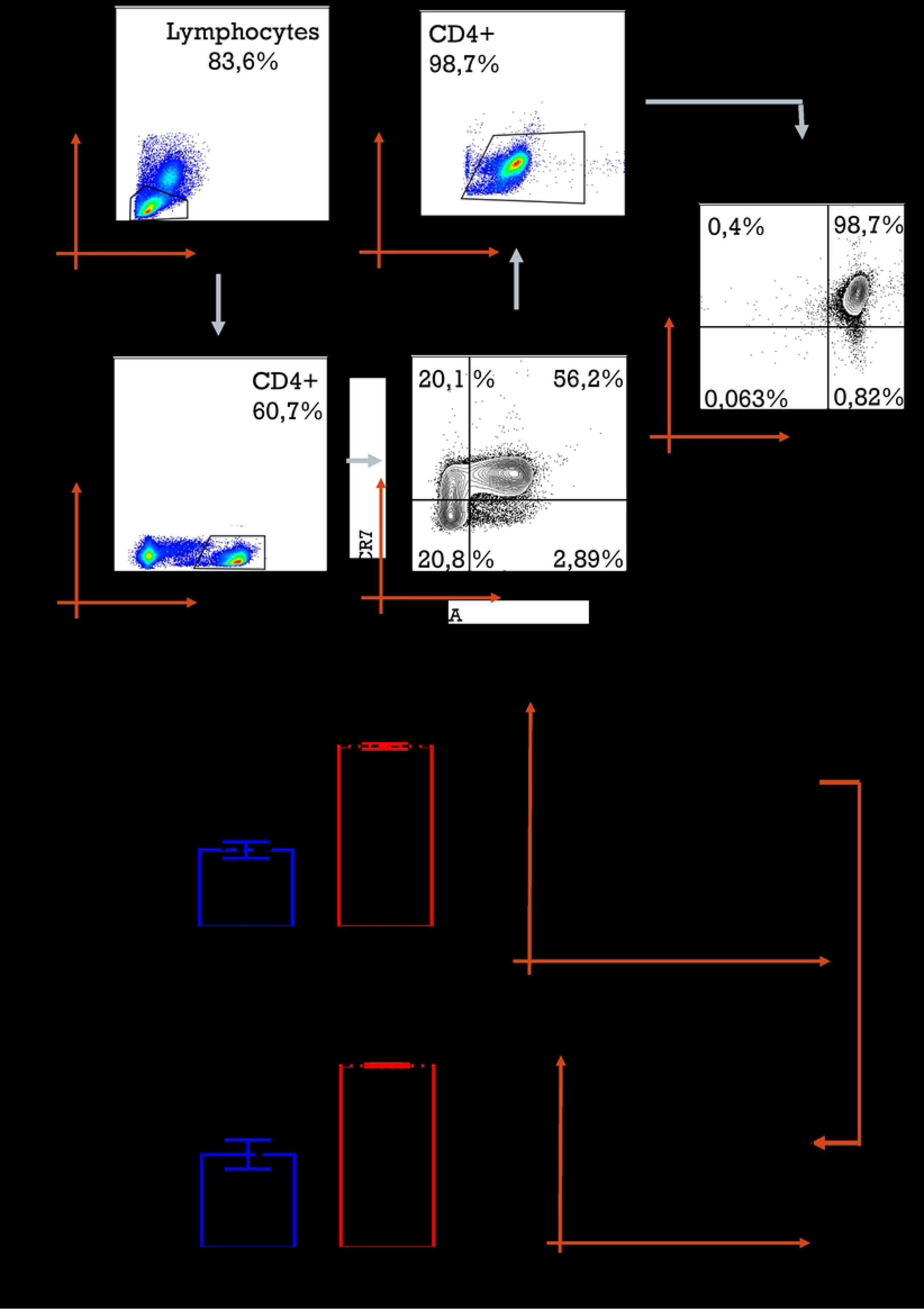
Gating strategy for enrichment of the naïve fraction of CD4 T cells and determining memory subsets. The purity of the naïve fraction enrichment was measured before and after magnetic separation by negative selection. In **A.** the gating strategy to define the naive component is shown. For all cultures, a purity >95% was obtained for CD4, and >97% for the naïve (CCR7^+^CD45RA^+^). n=10. Wilcoxon test. The bars correspond to the average + SEM. **= 0.0039. To determine the memory stem population, the strategy in **B.** was used: from the naive population (CCR7^+^CD45RA^+^), two subpopulations were obtained according to their CD95 expression, one CD95^−^ “true naive”, and one CD95^+^ corresponding to the memory stems (SCM). Representative plots of CD95 expression in the CCR7^+^CD45RA^+^ component at day 0 and day 15 of the cultures are shown in **B.** and **C.**, respectively.

**S2 Figure.**
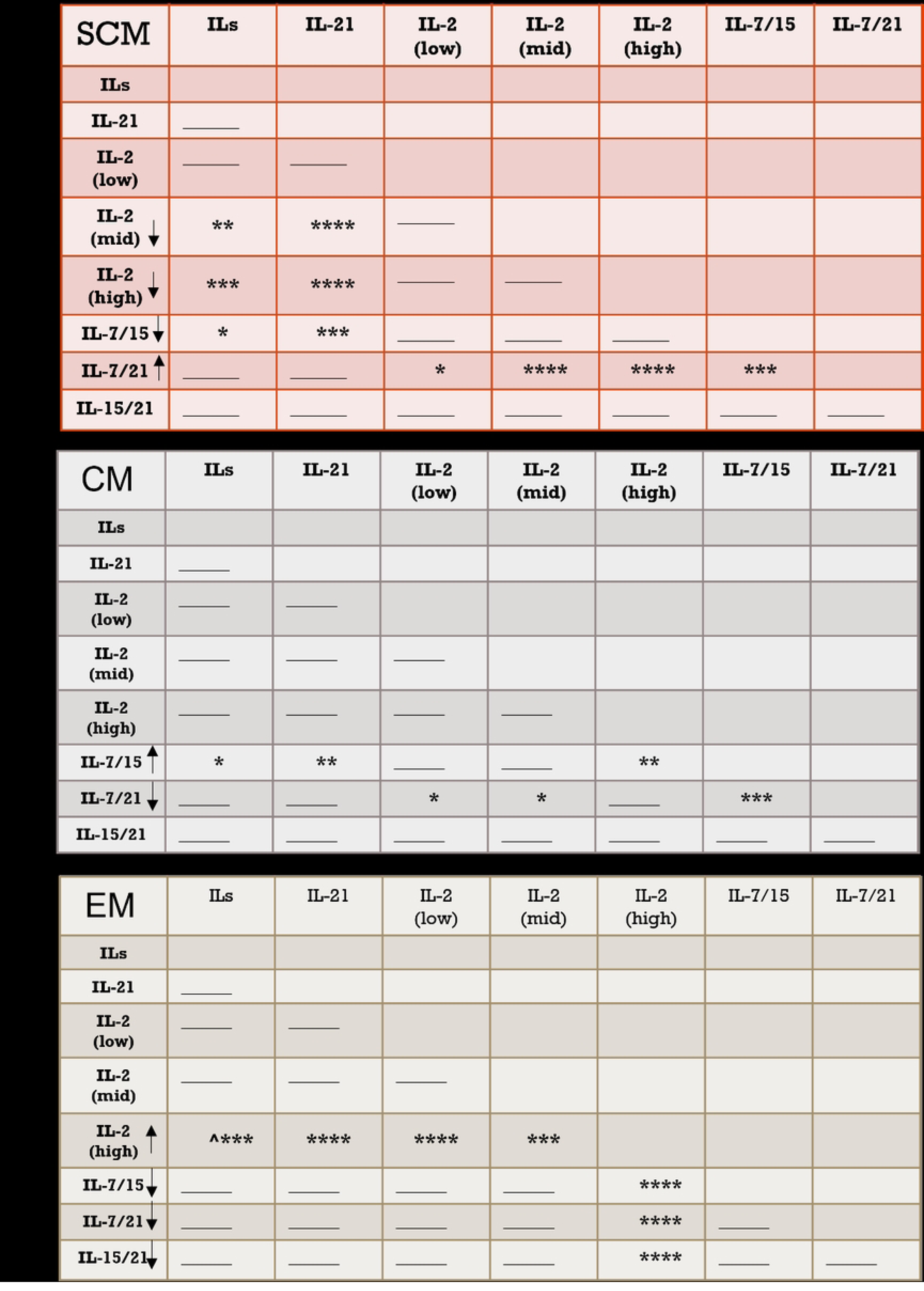
Memory profile on day 15 of the cultures. Two-way ANOVA of percentages of subpopulations at day 15 **A.** Stem memory (SCM), **B.** Central memory (CM) and **C.** Effector memory (EM) between the different treatments. 2.5 x10^5^ naïve CD4 T cells were stimulated for 15 days with anti-CD2/CD3/CD28 beads in a 2:1 lymphocyte-bead ratio accompanied with IL-7, 15 and 21; IL-21; IL-2 low; IL-2mid and IL-2 high; IL-7 and 15, IL-7 and 21, IL-15 and 21. n= 8. Non-significant changes are represented with a line.

**S3 Figure.**
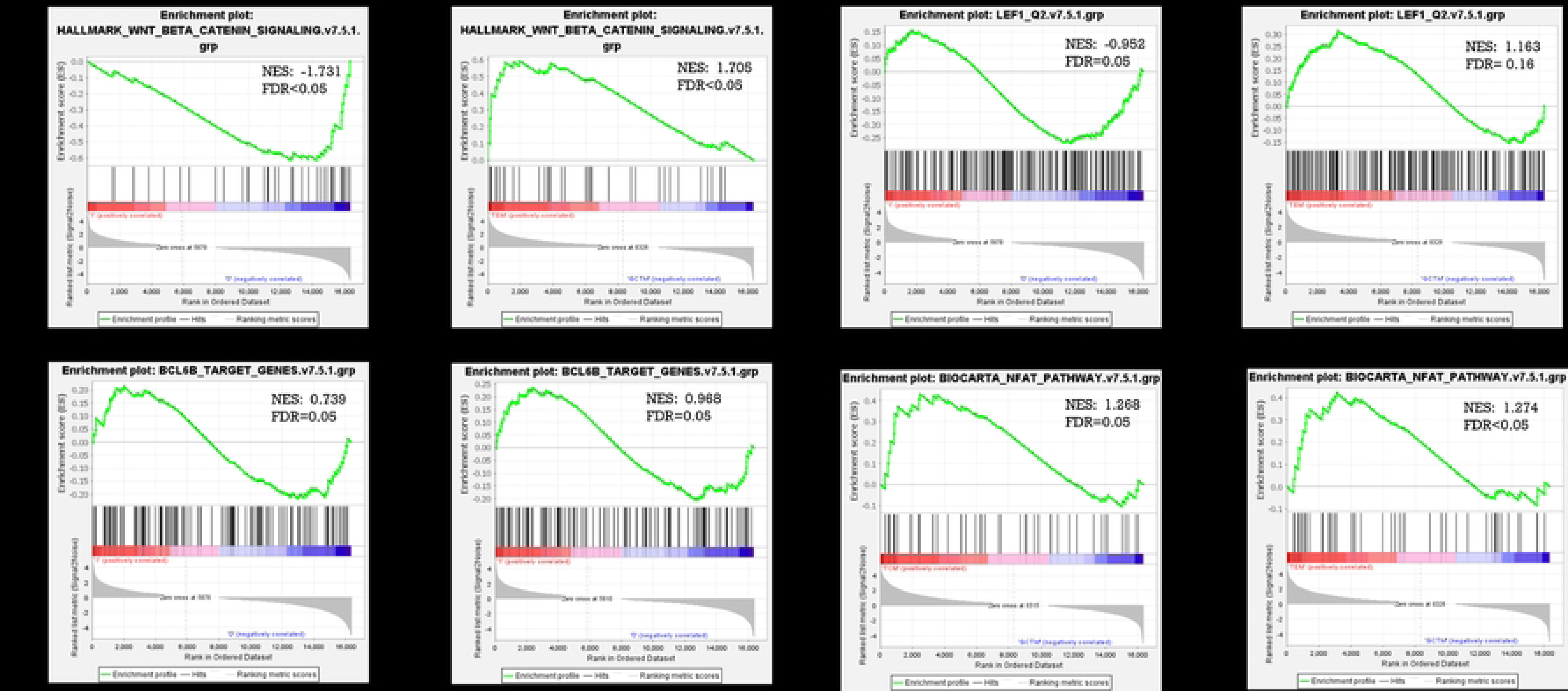
Analysis Enrichment of genes associated with self-renewal and activation pathways of CD4 T cells. RNA-seq of Tcm (GSE143215), Tfh (GSE130793) and Tem (GSE97863) cells were obtained from public databases and were normalized for analysis by GSEA. Gene pathways were obtained from the GSEA page on previously published pathways. GSEA-reported gene set enrichment plots for Wnt-β-catenin (M5895) between **A.** Tfh (red) vs Tcm (blue) and **B.** Tfh (red) vs Tem (blue), for Lef1 (M3572) between **C.** Tfh (red) vs Tcm (blue) and **D.** Tfh (red) vs Tem (blue), for Bcl-6 (M29904) between **E.** Tfh (red) vs Tcm (blue) and **F.** Tfh (red) vs Tem (blue), for NFAT (M2288) between **G.** Tfh (red) vs Tcm (blue) and **H.** Tfh (red) vs Tem (blue). The profile shows the continuous enrichment score (green curve) and the positions of gene set members (black vertical bars) in the rank-ordered list of differential gene expression. n= minimum 3.

**S4 Figure.**
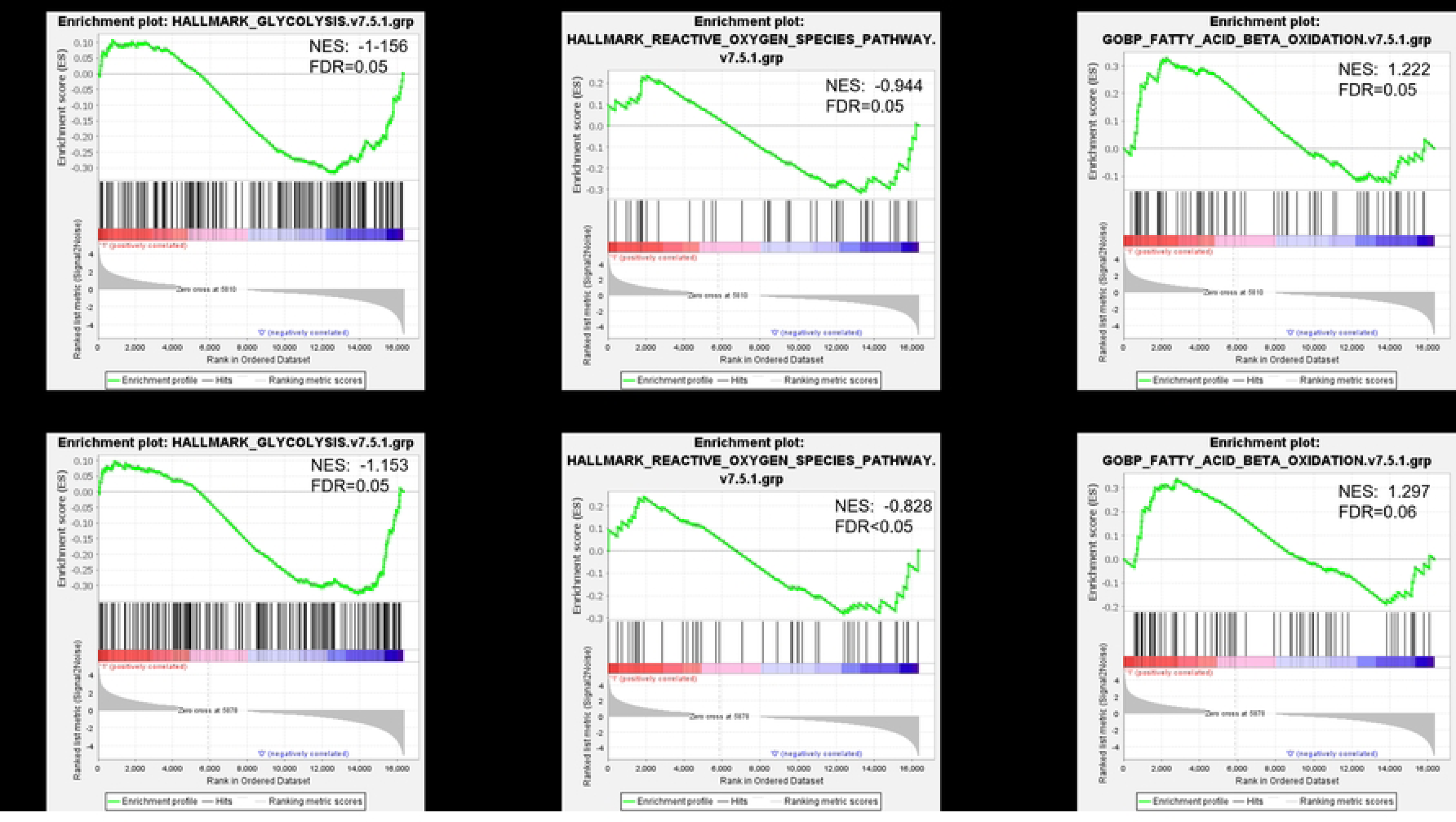
Analysis Enrichment of genes of pathways associated with metabolism. RNA-seq from Tscm (GSE143215), Tfh (GSE130793), and Tem (GSE97863) cells were obtained from public databases and normalized for analysis by GSEA. Gene pathways were obtained from the GSEA page on previously published pathways. Gene set enrichment plots reported by GSEA between Tfh and Tem for **A.** Glycolysis and **B.** ROS, and **C.** FAO and between Tfh vs Tcm for **D.** Glycolysis and **E.** ROS, and **F.** FAO. The profile shows the continuous enrichment score (green curve) and the positions of gene set members (black vertical bars) in the rank-ordered list of differential gene expression. n= minimum 3.

**S1 File. GSEA data of Tfh, Tscm, Tcm, and Tem subset.** GSEA data from each of the pathways included for the cell subsets comparisons.

